# Haplotype-phased common marmoset embryonic stem cells for genome editing using CRISPR/Cas9

**DOI:** 10.1101/373886

**Authors:** Bo Zhou, Steve S. Ho, Louis C. Leung, Thomas R. Ward, Marcus Ho, Melanie J. Plastini, Scott C. Vermilyea, Marina E. Emborg, Thaddeus G. Golos, Megan A. Albertelli, Philippe Mourrain, Dimitri Perrin, Karen J. Parker, Alexander E. Urban

**Affiliations:** Department of Psychiatry and Behavioral Sciences, Stanford University School of Medicine, Stanford, CA 94305, USA; Department of Genetics, Stanford University School of Medicine, Stanford, California 94305, USA; Stanford Center for Sleep Sciences and Medicine, Stanford University School of Medicine, Stanford, CA 94305, USA; Wisconsin National Primate Research Center, University of Wisconsin-Madison, Madison, WI 53715, USA; Neuroscience Training Program, University of Wisconsin-Madison, Madison, WI 53705, USA; Department of Medical Physics, University of Wisconsin-Madison, Madison, WI 53705, USA; Departments of Comparative Biosciences and Obstetrics and Gynecology, University of Wisconsin-Madison, Madison, WI 53715, USA; Department of Comparative Medicine, Stanford University School of Medicine, Stanford, California 94305, USA; Science and Engineering Faculty, Queensland University of Technology, Brisbane, QLD 4001, Australia; Program on Genetics of Brain Function, Stanford Center for Genomics and Personalized Medicine, Tasha and John Morgridge Faculty Scholar, Stanford Child Health Research Institute, Stanford University, Stanford, CA, USA

**Keywords:** *Callithrix jacchus*, transgenics, embryonic stem cells, haplotype phasing, CRISPR/Cas9, model organisms, marmoset, induced neurons (iNs)

## Abstract

Due to anatomical and physiological similarities to humans, the common marmoset (*Callithrix jacchus*) is an ideal organism for the study human diseases. Researchers are currently leveraging genome-editing technologies such as CRISPR/Cas9 to genetically engineer marmosets for the *in vivo* biomedical modeling of human neuropsychiatric and neurodegenerative diseases. The genome characterization of these cell lines greatly reinforces these transgenic efforts. It also provides the genomic contexts required for the accurate interpretation of functional genomics data. We performed haplotype-resolved whole-genome characterization for marmoset ESC line cj367 from the Wisconsin National Primate Research Center. This is the first haplotype-resolved analysis of a marmoset genome and the first whole-genome characterization of any marmoset ESC line. We identified and phased single-nucleotide variants (SNVs) and Indels across the genome. By leveraging this haplotype information, we then compiled a list of cj367 ESC allele-specific CRISPR targeting sites. Furthermore, we demonstrated successful Cas9 Endonuclease Dead (dCas9) expression and targeted localization in cj367 as well as sustained pluripotency after dCas9 transfection by teratoma assay. Lastly, we show that these ESCs can be directly induced into functional neurons in a rapid, single-step process. Our study provides a valuable set of genomic resources for primate transgenics in this post-genome era.

## Introduction

Biomedical research has been propelled by the ability to genetically modify conventional laboratory animals such as mice, fish, flies, and worms that have traditionally served as model organisms. These species, however, are evolutionarily distant to, and physiologically distinct from, humans, and are ill-suited for modeling complex brain functions and neuropsychiatric illness (Camus et al., 2015; Nelson and Winslow, 2009; Phillips et al., 2014). Although there is a critical need for developing sophisticated animal models with greater homology to human disease (Capitanio and Emborg, 2008; Emborg, 2017; Parker et al., 2018), until recently, transgenic non-human primate (NHP) research endeavors were a prohibitively inefficient and expensive undertaking (Grow et al., 2016). Recent breakthroughs in NHP genome-editing technologies, however, have advanced the possibility of gaining mechanistic insights into the genetic basis of human brain development and function.

The common marmoset (*Callithrix jacchus*) is an ideal model organism by which to advance the NHP engineering objective. Common marmosets are small New World monkeys and thrive in captivity under standard laboratory settings. Like humans, common marmosets are highly social, raise young biparentally, and demonstrate complex cognitive abilities (Ausderau et al., 2017; Schultz-Darken et al., 2016). Their metabolic, immunological, endocrinological, and anatomical characteristics are likewise quite similar to those of humans (Sasaki, 2015). Importantly, marmosets and humans share conserved brain structures, such as enlarged cerebral neocortices (Izpisua-Belmonte et al., 2015; Wise, 2008), as well as complex neural circuitry (Okano et al. 2016; Okano et al. 2012), rendering marmosets a well-suited model organism for the development of *in vivo* biomedical modeling for human neurological, psychiatric, and neurodegenerative diseases. Finally, marmosets also have inherent advantages over other NHPs, such as macaques, due to their small size, high reproductive efficiency, year-round ovarian cycles, ease of handling, and absence of serious zoonotic viral infections which can be fatal to humans (Mansfield, 2003).

Several marmoset embryonic stem cell (ESC) lines and induced pluripotent stem cell (iPSC) lines with the capabilities of differentiating into specified cell lineages are now available (Sasaki et al., 2005; Thomson et al., 1996; Tomioka et al., 2010; Vermilyea et al., 2017). The first transgenic primate showing successful germline transmission of transgenes was accomplished using the marmoset in which lentiviruses expressing enhanced green fluorescent protein (EGFP) were injected into preimplantation embryos (Sasaki et al., 2009). A similar technology was also used to generate transgenic marmosets expressing genetically encoded calcium indicators (Park et al., 2016; Sasaki et al., 2009; Tomioka et al., 2017). Shortly after the reporting of the first genetically modified marmosets, the marmoset genome was sequenced and assembled using various strategies including Sanger sequencing at 6X coverage (Marmoset Genome Sequencing and Analysis Consortium, 2014; Sato et al., 2015). Sequencing of the marmoset transcriptome across various tissues has also been undertaken in an effort to provide high-quality gene annotations for its recent genome assembly (Maudhoo et al., 2014). Furthermore, gene-editing technologies based on homologous recombination and programmable nucleases, such as zinc-finger nucleases (ZFNs) and transcription activator-like effector nucleases (TALENs), have been successfully applied in marmoset ESCs and embryos (Sato et al., 2016; Shiozawa et al., 2011).

Advances in CRISPR/Cas9 technology have enabled greater possibilities for NHP transgenic research including the introduction of copy number variations (CNVs) in animal models. The ability to create transgenic NHP animals using CRISPR/Cas9 was successfully demonstrated in cynomolgus monkeys in which multiple genes were altered in a single step (Niu et al., 2014). Efforts to engineer marmoset cells using CRISPR/Cas9 are currently underway in multiple labs around the world using the already-established marmoset ESC and iPSC lines (Vermilyea et al., 2018), and advances are expected in the coming years (Izpisua-Belmonte et al., 2015; Sato and Sasaki, 2018).

Genomic editing strategies stand to benefit from characterizing the genome of these cell lines. Whole genome characterization can assist the design of CRISPR gRNAs to gain sequence specificity and to decrease off target effects. Whole genome characterization also provides genomic contexts for accurate interpretation of functional genomics studies, such as measuring genome-wide expression levels and methylation states. Haplotype phasing across the genome allows for the identification of CRISPR sites suitable for allele-specific targeting. Whole-genome characterization in the context of copy-number variation (CNV) analysis also allows for the identification of large CNVs not evident from standard karyotyping analysis. This is especially important for studies that utilize *in vitro* differentiation of neurons, as they are known to harbor mosaic ploidy changes during development via a yet to be understood mechanism (Cai et al., 2014; Knouse et al., 2016; McConnell et al., 2013). Lastly, whole-genome characterization provides a reference genetic background for the transgenic animals or other cell lines derived from the ESCs.

Here, we performed whole-genome characterization in a haplotype-resolved manner for marmoset ESC line cj367 from the Wisconsin National Primate Research Center (Vermilyea et al., 2017). To the best of our knowledge, this is the first haplotype-resolved analysis of a marmoset genome and the first whole-genome characterization of any marmoset ESC line. Specifically, we identified single-nucleotide variants (SNVs) and Indels by performing deep-coverage whole-genome sequencing. We then phased these SNV and Indel haplotypes by performing linked-read sequencing at 128.5X physical coverage. Leveraging these phased haplotypes, we then compiled a list of cj367 ESC allele-specific CRISPR targeting sites. Furthermore, we demonstrated successful Cas9 Endonuclease Dead (dCas9) expression and targeted localization in cj367 as well as sustained pluripotency after dCas9 transfection by teratoma assay. We also performed independent confirmation of cj367 ESC pluripotency and also show that these ESCs can be directly induced into functional neurons in a rapid, single-step process. Our datasets provide a valuable set of resources for primate transgenic research.

## Results

### Characteristics of marmoset ESC line cj367 and direct induction to neurons

Characteristics of the cj367 ESCs were similar to that of human iPSCs, growing in dense, circular colonies adhered to the plate (Figure 1). Following the thawing and first passaging of the ESCs after arrival from the Wisconsin National Primate Research Center, removal of differentiated cells was only needed once. The ESCs required a daily change of media and to be passaged every 4 to 5 days (∼50-80% confluency). ESC colonies have clearly defined outlines, identical optical properties, and clear nuclei (Figure 1a). Extra care is needed when dissociating the cells from the plate because they readily detach with little to no agitation after adding a cell dissociation agent such as Accutase. Instead of aspirating the Accutase before adding PBS to fully dissociate the cells, DMEM/F12 media was added directly to the Accutase to wash the cells and then spun down to remove the Accutase/media mixture. The cj367 ESCs were also confirmed to be pluripotent by immunostaining for pluripotency markers NANOG, SSEA4, and OCT3/4 (Figure 1d-f) as was initially demonstrated elsewhere (Vermilyea et al., 2017).

**Figure 1.**
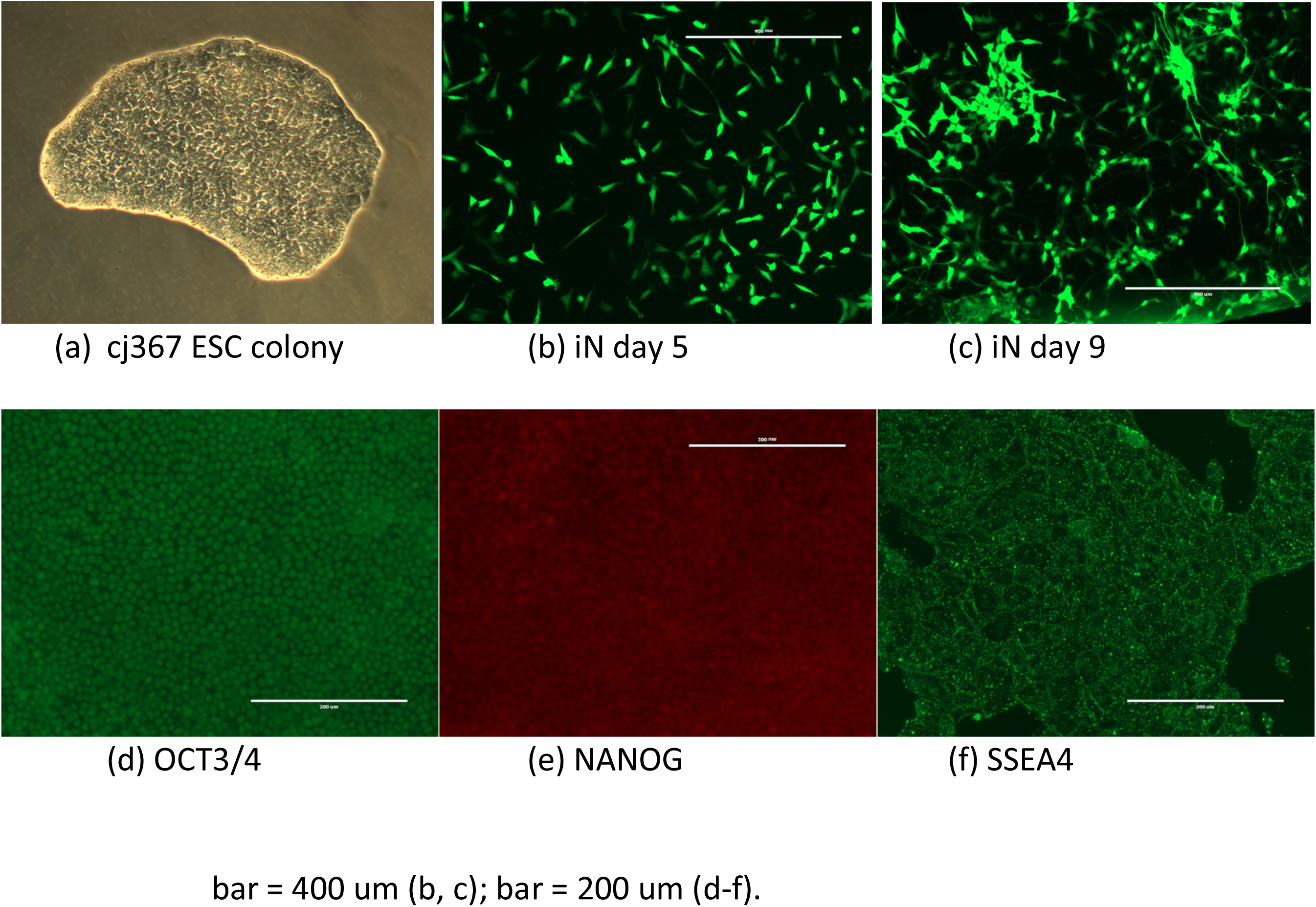
(a) cj367 ESC colony at 20X magnification. Days 5 (b) and 9 (c) respectively of cj367 directly induced into immature cortical glutamatergic neurons; white bar = 400 microns. Cells are marked by EGFP. Immunostaining of pluripotency markers OCT3/4 (d), NANOG (e), and SSEA4 (f) in cj367; white bar = 200 microns.

We differentiated the ESCs into immature cortical glutaminergic neurons using a two-week protocol originally designed and optimized for use in human iPSC lines (Zhang et al., 2013). To test the effectiveness of this protocol, the ESCs were plated at different densities based on total cell count (100k, 150k, 200k, 250k cells). Each well of cells received the same amount of the three lentiviral vectors described in (Zhang et al., 2013), and the protocol was followed for the next nine days. Cell density did not alter the results of the differentiation; however, the ESCs behaved differently from human iPSCs in two major ways. First, compared to human iPSCs, the cells that survived the differentiation process grouped entirely around the edge of the well, and only a few cells were observed in the middle. Second, the marmoset ESCs required 3 to 4 more days to reach complete differentiation compared to human iPSCs. The protocol was stopped short at day 9/10 for the marmoset ESCs, i.e. the human day-seven-equivalent (Zhang et al., 2013), and GFP fluorescence was used as a marker to visualize the differentiation process (Figure 1b, c).

### Identification of SNVs and Indels

From deep-coverage (∼60X) short-insert Illumina WGS of the cj367 genome, we identified SNVs and indels using GATK Haplotypecaller (McKenna et al., 2010). In standard GATK Best Practices workflow (DePristo et al., 2011) for human genomes, high-confidence SNVs and indels are filtered through a statistical learning approach using training datasets of variants derived from population-scale data from the 1000 Genomes Project (The 1000 Genomes Project Consortium et al., 2010) or the International HapMap Project (International HapMap Consortium, 2005). Since similar datasets are not available for marmosets, to identify high-confidence SNVs and indels, we adopted our own statistical approach. Briefly, we analyzed the distributions of the following parameters (QD, FS, MQ, MQRankSum, ReadPosRankSum) in the raw outputs from GATK Haplotypecaller and determined a series of appropriate filtering thresholds for SNVs and indels separately (see Materials and Methods).

Through this approach, we identified a total of ∼3.75M SNVs (2.23M heterozygous, 1.52M homozygous) and 0.82M indels (0.43M heterozygous, 0.38M homozygous) (Table 1, Figure 2a, EVA accession PRJEB27676); 33% of SNVs and 35% of indels are intragenic, and 0.73% of SNVs and 0.41% of indels are within protein coding exons (Table 1). Furthermore, we also identified long genomic stretches exhibiting continuous homozygosity by using a Hidden Markov Model (HMM) adopted from Adey and colleagues (Adey et al., 2013). In cj367, continuous stretches of chromosome 1, 2, 4, 6, 7, 9, 12, 13, 14, 15, and 22 show long continuous stretches of homozygosity that likely resulted from inbreeding (Figure 2a, Table S1).

**Figure 2.**
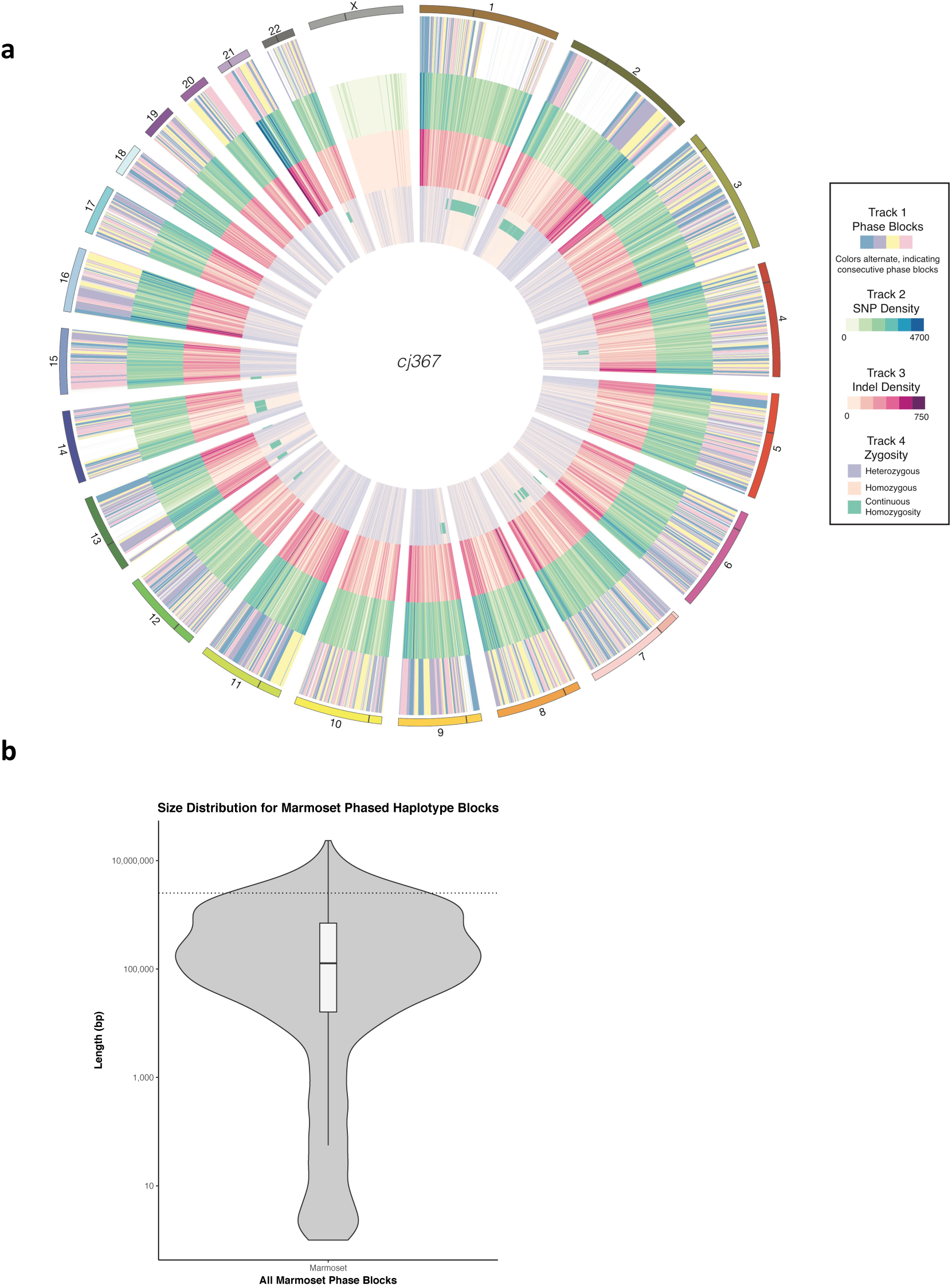
(a) Circos visualization of cj367 genome variants with the following tracks: 1. phased haplotype blocks (demarcated with 4 colors for clearer visualization); 2. SNV density in 1 Mb windows; 3. Indel density in 1 Mb windows; 4. zygosity (heterozygous or homozygous > 50%) in 1 Mb windows; regions with continuous homozygosity. (b) Violin plot of size distribution of haplotype blocks in cj367.

### Haplotype Phasing

We then haplotype-phased the identified SNVs and indels in cj367 by performing 10X Genomics Chromium linked-read library preparation and sequencing (Marks et al., 2018; Zheng et al., 2016). Post sequencing analysis showed that approximately 1.26 ng or 381 genomic equivalents of high molecular weight (HMW) genomic DNA fragments (average size 36 kb, 66.2% > 20kb, 14.3% >100kb) were partitioned into approximately 1.27 million droplets and uniquely barcoded (16 bp) (Table 1). The linked-read library was sequenced (2×151 bp) to 30.4× genome coverage with half of the reads coming from HMW DNA molecules with at least 30 linked-reads (N50 Linked-Reads per Molecule) (Table 1). We estimate the actual physical coverage (C_F_) to be 128.5×. Coverage of the mean insert by sequencing (C_R_) is 8,520 bp (284bp × 30 linked-reads) or 23.7% of the average input DNA fragment size, thus the overall sequencing coverage C = C_R_ × C_F_ = 30.4×. In 3115 haplotype blocks (Table 1, EVA accession PRJEB27676), 2.16M (96.9%) of heterozygous SNVs and 0.79M (83.7%) of indels were phased. The longest phased haplotype block is 23.4 Mbp (N50 = 2.49Mbp) (Figure 2b, Table 1, EVA accession PRJEB27676). Haplotype block sizes vary widely across chromosomes (Figure S1, Figure 2a), and poorly phased regions correspond to regions exhibiting continuous homozygosity (Table S1, Figure 2a, EVA accession PRJEB27676).

### Cj367 genome-specific and allele-specific CRISPR targets

We identified 44,590 targets in cj367 genome suitable for allele-specific CRISPR targeting (Table S2). Phased variant sequences (including reverse complement) that differ by >1 bp between the alleles were extracted to identify all possible CRISPR targets by pattern searching for [G, C, or A]N_20_GG (see Materials and Methods). Only conserved high-quality targets were retained by using a selection method previously described and validated (Sunagawa et al., 2016). We took the high-quality target filtering process further by taking the gRNA function and structure into account. Targets with multiple exact matches, extreme GC fractions, and those with TTTT sequence (which might disrupt the secondary structure of gRNA) were removed. Furthermore, we used the Vienna RNA-fold package (Lorenz et al., 2011) to identify gRNA secondary structure and eliminated targets for which the stem loop structure for Cas9 recognition is not able to form (Nishimasu et al., 2014). Finally, we calculated the off-target risk score using the tool as described for this purpose (Ran et al., 2013). A very strict off-target threshold score was chosen in which candidates with a score below 75 were rejected to ensure that all targets are as reliable and as specific as possible.

### CRISPR/Cas9 in marmoset ESC

In order to test whether CRISPR/Cas9 technology would be suitable to create transgenic marmoset models, we tested whether sgRNA-guided CRISPR entry was possible in cj367. After transfecting cj367 marmoset ESCs with plasmids carrying *dCas9-GFP* (Chen et al., 2013) and using sgRNA sequences that target telomeres, we successfully located dCas9-GFP fluorescence from telomeres within the nuclei of live marmoset ESCs (three nuclei shown; Fig. 3a-c), demonstrating that the Cas9 nuclease can indeed enter the nuclease and possibly edit the genome in a targeted fashion. Teratoma assay demonstrate that pluripotency of ESC remains unchanged after dCas9-GFP transfection (Figure 4).

**Figure 3.**
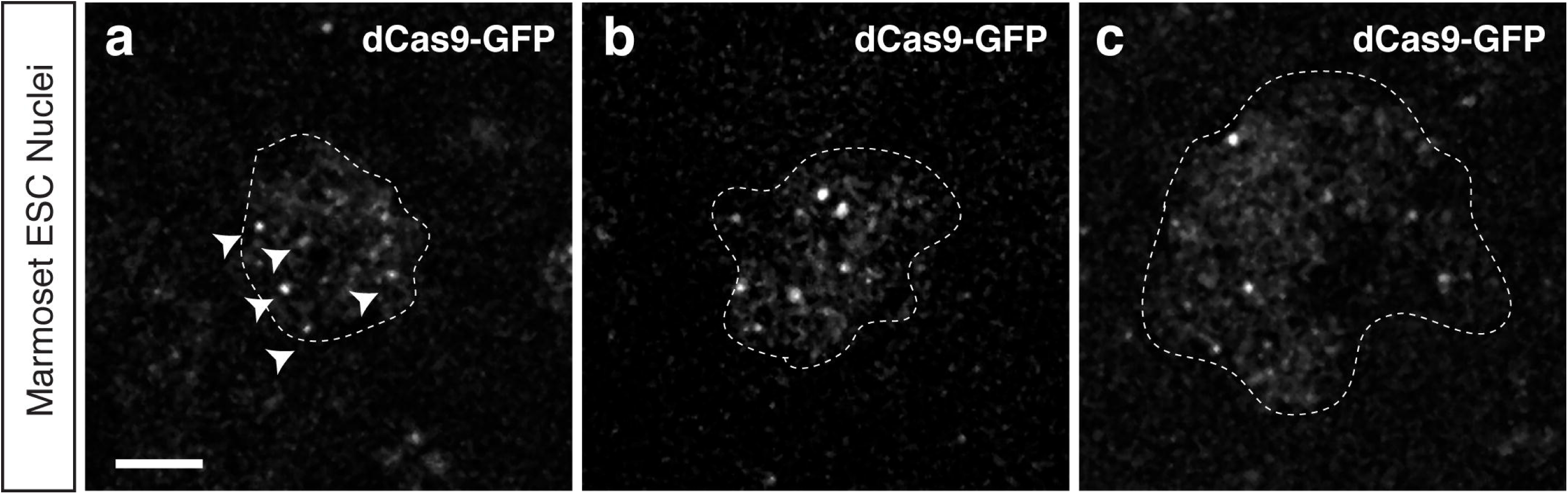
(a-c) Three examples of marmoset ESCs transfected with dCas9-GFP that are targeted telomeric sequences revealing telomeres as fluorescent puncta (white arrowheads). Images are maximum projections of the nuclear volume and the dashed outlines indicate the nuclear boundaries. Scale bar, 5 µm.

**Figure 4.**
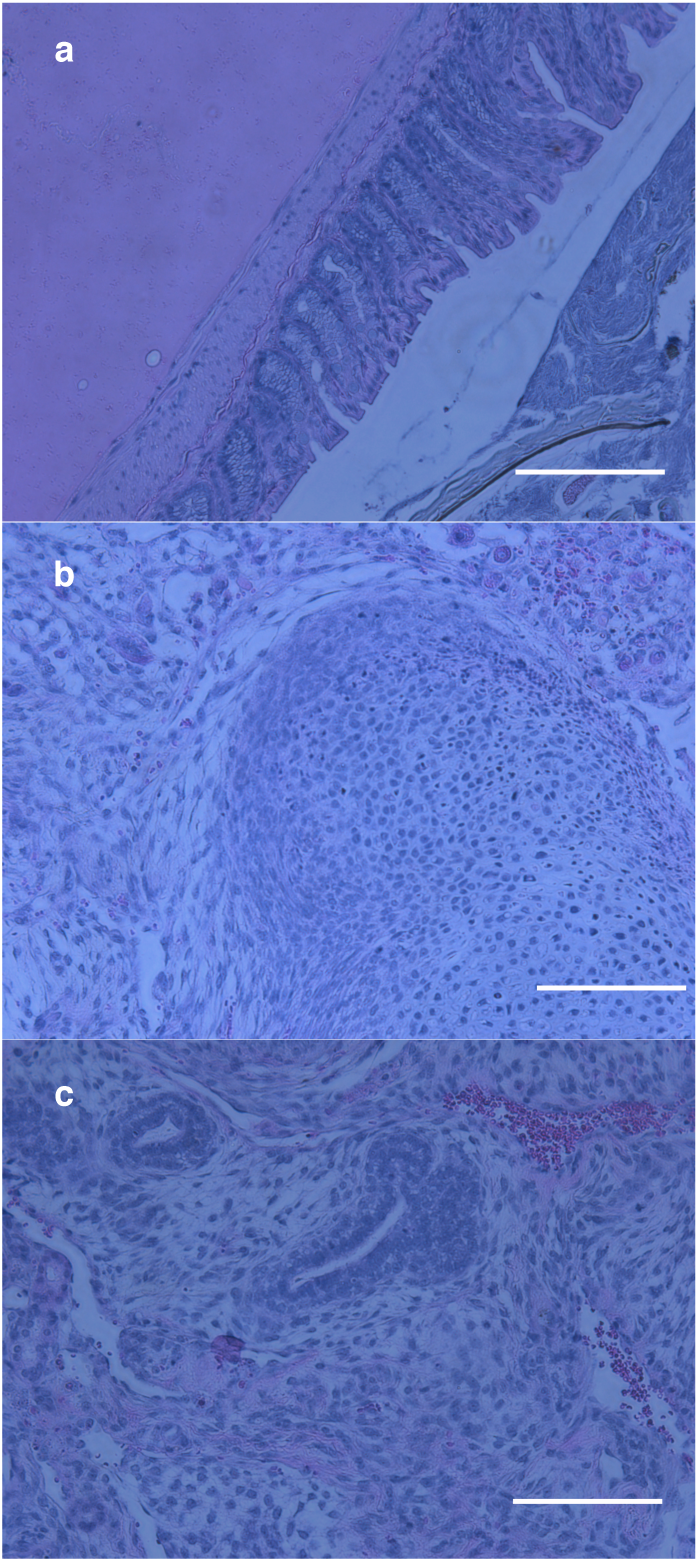
Histological analysis of teratomas derived from marmoset ESCs after dCas9-GFP transfection. Scale bar is 50 µm. Mamorset lines are characterized by pluripotency as demonstrated by their ability to form glandular epithelium (endoderm in a); cartilage (mesoderm in b), and neural rosettes-like structures (ectoderm in c).

## Discussion

One of the many advantages of using the common marmoset as a model to study human disease is that its physiology, brain structure, and aspects of behavior more closely resemble that of humans (Kishi et al., 2014; Okano et al., 2012). The creation of pluripotent marmoset SC lines provides a platform on which to study molecular and cellular patterns typical of the species. Traditionally, marmoset ESC lines have been derived from the inner cell mass of blastocysts, and all of the surviving lines had been female (Debowski et al., 2016; Sasaki et al., 2005; Thomson et al., 1996). Recently, three novel marmoset ESC lines have been derived from the natural morula stage of preimplantation embryos, and one novel male ESC line has been derived from the expanded blastocyst stage (Debowski et al., 2016). Marmoset iPSCs from fetal liver cells and adult fibroblasts have also been reported (Farnsworth et al., 2013; Qiu et al., 2015; Tomioka et al., 2010; Vermilyea et al., 2017). The capacity of marmoset ESC and iPSC lines to differentiate into cardiomyocytes and hematopoietic cells by induction has been demonstrated (Sasaki et al., 2005; Tomioka et al., 2010). They can also be directed to neural cell differentiation using the stromal cell-derived inducing activity method and to dopaminergic neurons using a dual-SMAD-inhibition induction method (Sasaki et al., 2005; Tomioka et al., 2010; Vermilyea et al., 2017). Here, we independently validate the pluripotency of the cj367 cell line (Figure d-f) and show that ESC line cj367 can be directly induced to neurons in a rapid, single step using a protocol developed for human pluripotent stem cells (Zhang et al., 2013) (Figure 1b, c). In addition to revealing these biological characteristics of cj367 that are especially applicable for neuroscience research, our results allow researchers to perform genome-editing in cj367 using CRISPR/Cas9 in a “personalized” and allele-specific manner. Our haplotype-resolved whole-genome characterization of cj367 also provides a reference for all future functional genomic and epigenomics studies, such as RNA-seq, DNA methylation analysis, or ChIP-seq, carried out using cj367 or derivative cell lines/animals. Taken together, our data provide a valuable resource for the primate transgenic research community in this post-genome era.

The recent successes in using CRISPR-Cas9 to disrupt specific genes such as *Ppar-*γ, *Rag1*, and *Dax1* in cynomolgus (Kang et al., 2015; Niu et al., 2014) and *DMD* in rhesus monkeys (Chen et al., 2015) forecast that the use of CRISPR-Cas9 to genetically edit marmoset cells or embryos is within reach (Izpisua-Belmonte et al., 2015; Kishi et al., 2014; Kropp et al., 2017; Sato and Sasaki, 2018). Gene-editing technologies that leverage programmable nucleases, such as ZFNs and TALENs have already been demonstrated in marmoset ESCs (Sato et al., 2016; Shiozawa et al., 2011). Here, we provide further evidence on this exciting possibility by demonstrating the successful sgRNA-guided CRISPR entry and expected dCas9-GFP localization in marmoset ESCs, which suggests that the Cas9 nuclease can indeed enter the nucleus and edit the genome in a sequence-directed manner. Furthermore, this not only demonstrates the relative ease of porting existing gene-editing methodologies but also the exciting future feasibility of creating genetic models of human disorders that provide new insights only possible in NHP models (Capitanio and Emborg, 2008; Zahs and Ashe, 2010).

With the imminent application of CRISPR-Cas9 for marmosets on the horizon, it is important to have genome characterizations of the NHP ESC and iPSC lines that are being distributed to various research labs across the world from primate cell repositories. It is now widely recognized that sequence variants can confound the intended on-target and off-target sites of the gRNAs especially when appropriate controls are lacking (Pinello et al., 2016). The CRISPR system relies on Watson and Crick base pairing to mediate genomic editing, thus it is not surprising that genetic sequence variation can widely affect its efficiency. By taking advantage of the extensive documentation of human genetic variation, efforts have been made to elucidate the impact of genetic variation on CRISPR targeting (Canver et al., 2018). Importantly, it has also been recognized that this effect on CRISPR-targeting efficiency due to genetic sequence variation can be leveraged to make cell line-specific or allele-specific CRISPR edits for various studies (Zhou et al., 2018). The need for such resources to facilitate the use of CRISPR-Cas9 in primate research has already been voiced in the literature (Luo et al., 2016). Here, we performed the first whole-genome characterization for an established NHP ESC line and the first haplotype-phasing for a marmoset genome. We also determined the CRISPR-targeting sites suitable for allele-specific editing for cj367 (Table S2). Our haplotype-resolved whole-genome analysis as well as a list of CRISPR-targeting sites will not only be useful for cj367 in particular but also for future cell lines and animal models derived from the same genome as cj367.

Possessing the unique property among mammals of routinely producing dizygotic twins that exchange hematopoietic stem cells *in utero*, all marmoset animals are naturally chimeric (Benirschke et al., 1962). While such chimerism will affect analyses of DNA samples obtained from developed animals, it does not affect our whole-genome of ESCs since the cj367 ESC line was derived before the developmental stage where the differentiation and exchange of hematopoietic stem cells can occur.

Another important aspect of whole-genome characterization is the analyses of structural and copy-number variation. The current quality of the marmoset reference genome assembly does not allow for confident mapping of structural variants (Marmoset Genome Sequencing and Analysis Consortium, 2014; Sato et al., 2015). Improvements in this regard are greatly needed. This also applies to other NHP genome assemblies as well (Luo et al., 2016).

Our whole-genome analysis of marmoset ESC line cj367 also allows for the accurate and confident interpretation of functional genomics datasets. Recently, RNA-seq experiments have been carried out in marmoset ESCs that show that overall expression patterns differ between cell lines derived from different animals under the same culturing conditions (Debowski et al., 2016), which could be a result of different genetic backgrounds. Furthermore, the expression patterns of various marmoset ESC lines more closely resemble primed pluripotency states rather than naïve states even though these cells passed all criteria for expression of established pluripotent markers (Debowski et al., 2016). This suggests that the culturing methods established for human pluripotent cells are not directly applicable to marmoset cells. Importantly, these results also suggest that functional genomic and epigenomics studies such as RNA-seq, ATAC-seq, and whole-genome bisulfite sequencing could be adopted as a tool to screen for and identify marmoset ESC lines that may exhibit true naïve pluripotency states.

## Materials and Methods

### Cell Culture

The frozen marmoset ESC starter cultures were removed from the liquid nitrogen tank (−80 °C) and placed in a 37 °C water bath until thawed. Cells were transferred to a 15 mL centrifuge tube and 10 mL DMEM/F12 (Thermo Scientific, SH30023.01) was added drop-wise in order to minimize bubble formation. The mixture was centrifuged at 1000 RPM for 3 min. The supernatant was aspirated and the process was repeated allowing the cells to be washed once more with another 10 mL DMEM/F12. Cells were re-suspended in 1 mL of medium, which was added to a single well of Matrigel hESC-qualified Matrix (BD Biosciences, 356234)-coated BioLite 6-well clear multi-dish (Thermo Scientific, 130184) and incubated at 37 °C and 5% CO_2_. The medium in each well was changed daily and the cells were passaged every 4 to 5 days. For passaging, cells were washed with DPBS (Life Technologies, 14190-144), dissociated with Accutase (Stem Cell Technologies, 7920), and added to fresh medium on a new plate and/or added to freezing medium for storage.

### Media ingredients

- E8 Medium (Thermo Scientific, A15169-01)
- E8 Supplement (Thermo Scientific, A15171-01)
- Glutamax (Life Technologies, 35050-061)
- Lipid Concentrate (Life Technologies, 11905-031)
- Nodal (R&D Systems, 3218-ND)
- Glutathione (Sigma, G4251)

Freezing medium: 20% DMSO (Thermo Fischer, 20688) solution (DMSO + DMEM/F12).

### Differentiating marmoset ESCs into iNs

When the marmoset ESCs reached confluence, the cells were dissociated using Accutase and plated at 250,000, 200,000, 150,000, and 100,000 cells per well. One-day post-passage, a single well was transfected with the 3 viral vectors as described in the two-week iN protocol (Zhang et al., 2013). The vectors were acquired from the Genome Virus and Vector Core (GVVC) at Stanford Neuroscience Institute (neuroscience.stanford.edu). The protocol was stopped prior to plating on mouse glia in order to achieve immature cortical glutamatergic neurons. Following transfection, cells were subjected to puromycin and doxycyclin selection and then monitored for a minimum of 7 days. For human iPSCs, full differentiation is usually completed at day 7. The marmoset ESCs required 10 to 11 days (the day of passage is defined as “day 0”) to achieve full differentiation.

### CRISPR-Cas9 Transfection

Marmoset cj367 embryonic stem cells were seeded onto a 35mm glass bottom microwell dish (MatTek) that was coated with 0.083 μg of BD Matrigel (BD Biosciences) per well. While the cells were maintained for approximately 48 hours to recover and return to their typical growth rate, they were visually scanned under a light microscope and manually dissected to remove any patches of spontaneously differentiated cells. At 70% confluence on the 35mm dish, the cells were transfected simultaneously with Addgene plasmids pSLQ1658-dCas9-EGFP and pSLQ1651-sgTelomere(F+E). We followed Invitrogen’s protocol for the Lipofectamine LTX transfection reagent (Thermo Fisher Scientific) using 5μL with 1μg of DNA from each plasmid for one 35mm dish. The lipid-DNA complexes were adjusted to a final volume of 100μL using OptiMEM (Thermo Fisher Scientific) and added to the marmoset cells for six hours. Following six hours of incubation, medium was changed to its original E8 formula. Five hours before imaging, cells were incubated with a photostable DMEM (MEMO EMD Millipore) supplemented with ROCK inhibitor to reduce photobleaching effects of the E8 media. Finally, an additional medium change using MEMO was done immediately before imaging.

### Nuclear imaging

Marmoset cell nuclei were imaged with an OMX BLAZE 3D-Structured Illumination Microscope (Applied Precision, GE). Cells were transferred to a modified stage adaptor held at 37°C and 5% CO_2_. 1μm z-stacks (0.125μm intervals) were taken at 100x magnification (U-PLANAPO 100X SIM objective N.A. 1.42) with a 488nm laser using DeltaVision OMX acquisition software (version 3.70.9622.0, GE) and deconvolved with SoftWoRx 7.0.0 software (Deltavision) before maximum projections were made for the figures (Fiji/ImageJ).

### Teratoma Generation and Histopathology

H9 (2.5million/5ul matrigel mixed with 25ul of PBS) and marmoset ES cells (3.5million/5ul matrigel mixed with 30ul of PBS) were injected subcutaneously or into the kidney capsule in 8-week-old male NSG mice (Jackson). Seven weeks after injection, the mice were euthanized and the teratomas were harvested. All animal studies were approved by Stanford University IACUC guidelines. For histological analysis, slides were stained with hematoxylin and eosin (H&E).

### Illumina WGS

Marmoset ESC (cj367) genomic DNA was extracted using the Qiagen DNeasy Blood & Tissue Kit (Cat.D No. 69504) and quantified using the Qubit dsDNA HS Assay Kit (Invitrogen, Waltham, MA, USA). DNA purity was verified (OD260/280 > 1.8; OD260/230 > 1.5) using NanoDrop (Thermo Scientific, Waltham, MA, USA). Afterwards, DNA was sheared to 200 bp to 850 bp (average size = 400 bp) using the Covaris S2 instrument (Covaris, Woburn, MA, USA). DNA was then concentrated to a volume of 50 µL using the DNA Clean & Concentrator kit (Cat. No. D4013) from Zymo Research (Irvine, CA, USA). One short-insert WGS library was constructed using the Kapa Hyper Prep Kit (KR0961) from Kapa Biosystems (Woburn, MA, USA) with 150 ng of sheared DNA as input following standard manufacturer’s protocol with 10 cycles of PCR. Library-size selection (420 bp to 1 kb) was carried out on the BluePippin instrument using the R2 marker on a 1.5% dye-free cassette from Sage Science (Beverly, MA, USA). WGS library-size selection was then verified using the 2100 Bioanalyzer DNA 1000 Kit (Agilent Technologies, Santa Clara, CA, USA). The WGS library was then sequenced (2×150 bp) on two lanes of the Illumina HiSeq 4000 to achieve >600 million pass-filter paired-end reads.

### Determining SNVs and Indels

Paired-end reads from the two lanes of Illumina HiSeq 4000 were combined and aligned to the *Callithrix jacchus* genome (GenBank accession GCA_000004665.1) using BWA-MEM version 0.7.5 (Li and Durbin, 2009) followed by marking of duplicates using Picard tools (version 1.129) (http://broadinstitute.github.io/picard/). SNVs and Indels were called using by GATK Haplotypecaller (version 3.7) (McKenna et al., 2010) and hard filtered for quality (QD > 2.0, FS < 60.0, MQ > 40.0, MQRankSum > −12.5, ReadPosRankSum > −8.0).

### Identifying genomic regions exhibiting continuous homozygosity

A Hidden Markov Model (HMM) was used to identify genomic regions exhibiting continuous homozygosity. The HMM is designed with two states: homogzyous and heterozygous. We used the hard-filtered SNVs we derived from GATK HaplotypeCaller, and filtered again for GQ > 30 and QUAL > 50. The genome was split into 25 kb bins; heterozygous and homozygous SNVs were tallied for each bin, and bins with <25 SNVs were removed. A bin was classified as heterozygous if >50% of the SNVs within the bin are heterozygous, otherwise it was classified as homozygous. This classification was used as the HMM emission sequence. The HMM was initialized with the same initiation and transition probabilities (Prob=10E-8) (Adey et al., 2013), the Viterbi algorithm was used to estimate a best path, and adjacent homozygous intervals were merged.

### 10X Genomics linked-read library and sequencing

Marmoset ESC (cj367) genomic DNA was extracted using the MagAttract HMW DNA Kit from Qiagen (Cat. No. 67563) (Hilden, Germany). The extracted genomic DNA was verified to be of HMW (average size >30 kb) using using field-inversion gel electrophoresis on the Pippin Pulse System (Sage Science, Beverly, MA, USA). DNA was then diluted to 1 ng/µl to be used as input for the 10x Genomics (Pleasanton, CA, USA) Chromium system (Marks et al., 2018; Zheng et al., 2016) in which HMW DNA fragments are partitioned into >1 million droplets in emulsion, uniquely barcoded (16 bp) within each droplet, and subjected to random priming and isothermal amplification following standard manufacturer’s protocol. The barcoded DNA molecules were then released from the droplets and converted to a linked-read library in which each library molecule has a 16 bp “HMW fragment barcode”. The final linked-read library (8 cycles of PCR amplification) was quantified using the Kapa Library Quantification Kit (Cat. No. KK4824) from Kapa Biosystems (Woburn, MA, USA) and diluted to 4 nM. Sequencing (2×151 bp) was performed on the Illumina NextSeq 500 using the NextSeq 500/550 High Output v2 kit (Cat. No. FC-404-2004) to achieve >30× genomic coverage.

### Haplotype phasing using linked-reads

Read-pairs i.e. linked-reads, that come from the same HMW DNA fragment carry the same “HMW fragment barcode” on Read 1. Virtual long-reads that are representative of the sequences of the original HMW genomic DNA fragments can be generated by clustering of linked-reads with identical barcodes. This process allows for the haplotype phasing of heterozygous SNVs and Indels. Paired-end linked-reads (median insert size 368 bp, duplication rate 5.39%, Q30 Read1 75.6%, Q30 Read2 68.4%) were aligned to *Callithrix jacchus* genome (GenBank accession GCA_000004665.1) (alignment rate 87.3%, mean coverage 30.4x, zero coverage 0.938%) and analyzed using the Long Ranger Software (version 2.1.3) from 10x Genomics (Marks et al., 2018; Zheng et al., 2016) (Pleasanton, CA, USA). Genome indexing was performed prior to alignment using the Long Ranger *mkref* module. Since Long Ranger *mkref* requires less than 500 contigs, we concatenated contigs in GCA_000004665.1 to meet this requirement. Reference gaps and repeat regions, downloaded from UCSC Genome Browser (Karolchik et al., 2004; Kuhn et al., 2013), were excluded from the analysis. Phasing was performed by specifying the set of pre-determined and filtered heterozygous SNVs and Indels from using GATK (see above) and formatted using *mkvcf* from Long Ranger (version 2.1.5). Phasing analysis was performed using the Long Ranger *wgs* module.

### Allele-specific CRISPR targets

To identify allele-specific CRISPR targets, we started by extracting variants that satisfy the following properties from Dataset S2 (phased variants):

1. They passed quality control (VCF field ‘Filter’ is equal to “PASS”)
2. They are phased (VCF field ‘GT’ uses “|” as separator rather than “/”)
3. The alleles are heterozygous (e.g. “0|1” or “1|0”, but not “1|1”)
4. The difference between the alleles is > 1 bp (to ensure target specificity) For the 704,571 variants that satisfy these four properties, we extracted the two haplotype sequences (maximum length: 595). We only worked with the sequences that were present in the phased genotype. Extracted sequences were tagged them according to their haplotype, for instance:

- If the sequence in the ‘Ref’ field was “GTA”, the ‘Alt’ field sequence was “TA”, and phasing was “0|1” i.e. Haplotype_1|Haplotype_2”, the sequence containing “GTA” was tagged “1” for Haplotype 1 and the sequence containing “TA” was tagged “2” for Haplotype 2.
- If the ‘Ref’ field was sequence “GTA”, the ‘Alt’ field was “TA,GCTA”, phasing was “1|2”, the sequence containing “TA” is tagged “1”, the sequence containing “GCTA” was tagged “2” (and the sequence containing “GTA” was not used)

A regular expression, RegEx, was used to extract all potential CRISPR targets from these sequences (i.e. all sequences that matched a [G, C, or A]N_20_GG pattern and those for which the reverse-complement matched this pattern). This yielded 720,239 candidates, which were then filtered to retain only high-quality targets. The process is adapted from a selection method previously described and validated (Sunagawa et al., 2016) and has already been used for more than 20 genes (Tatsuki et al., 2016). A high-quality candidate gRNA needs to target a unique site. All the candidates that have multiple exact matches in hg19 (irrespective of location) were identified using Bowtie2 (Langmead and Salzberg, 2012) and removed. We also removed targets with extreme GC content (>80% or <20%), and targets that contain TTTT, which tends to break the gRNA’s secondary structure. We also used the Vienna RNA-fold package (Lorenz et al., 2011) to compute the gRNA’s secondary structure. We eliminated all candidates for which the stem loop structure cannot fold correctly for Cas9 recognition (Nishimasu et al., 2014), except if the folding energy was above −18 (indicating that the “incorrectly-folded” structure is unstable). Finally, we evaluated the off-target risk score using our own implementation of the Zhang tool (Ran et al., 2013). To ensure that all targets are as reliable and specific as possible, we used a very strict threshold and rejected candidates with an off-target risk score less than 75. Candidates that satisfy all these requirements are considered high quality. For each candidate, we report the location of the variant (chromosome and position), the haplotype (“1” or “2” from the “Phased Genotype” column i.e. “Haplotype_1 | Haplotype_2”), the gRNA target sequence, its position relative to the start of the variant, its orientation, its off-target score, and the genomic element targeted (gene or enhancer). Note that the position relative to the start of the variant is for the 5’-most end of the target respective to the genome: if the target is 5’-3’, it is its 5’ end; if a target was extracted on the reverse complement, it is its 3’ end.

### Data access

All sequencing data are available via NCBI SRA accession SRP141443. Variant data is available via European Variation Archive accession PRJEB27676.

## Acknowledgements

This research was supported by a Tashia and John Morgridge Faculty Scholar Award from the Stanford Child Health Research Institute (AEU), a the Stanford Neurosciences Institute Seed Grant (AEU & KJP), the Stanford Child Health Research Institute Grant (AEU & KJP), and the Stanford Medicine Faculty Innovation Program (AEU). BZ was funded by NIH grant T32 HL110952. LCL was funded by the Stanford School of Medicine Dean’s Fellowship and NIH grants NS062798, DK090065, and MH099647. We are thankful for the support of the Stanford CSIF core facility – the imaging with the DeltaVision OMX system described was supported, in part, by award 1S10OD01227601 from the National Center for Research Resources (NCRR). This research was partially supported by NIH grants R24OD019803 to TGG and MEE, and P51OD011106 to the Wisconsin National Primate Research Center. The sequencing data was generated on an Illumina HiSeq 4000 that was purchased with funds from NIH award S10OD018220. Lastly, we thank Arineh Khechaduri for performing marmoset ESC genomic DNA extraction. We also thank Dr. Jeewon S. Kim and Dr. Vittorio Sebastiano at the Transgenic, Knockout and Tumor Model Center of the Stanford Cancer Institute for teratoma analyses.

## Disclosure Declaration

The authors of this manuscript declare no conflicts of interest.

## Author Contributions

B.Z., L.C.L., T.R.W., M.H., and M.J.P. performed experiments. B.Z., S.S.H., and D.P. performed genomic data analysis. S.C.V., M.E.E., T.G.G., P.M., K.J.P., and A.E.U. provided materials and reagents. K.J.P. and A.E.U. jointly supervised the study. B.Z. wrote the manuscript with suggestions from all co-authors.

---

**Figure S1.**
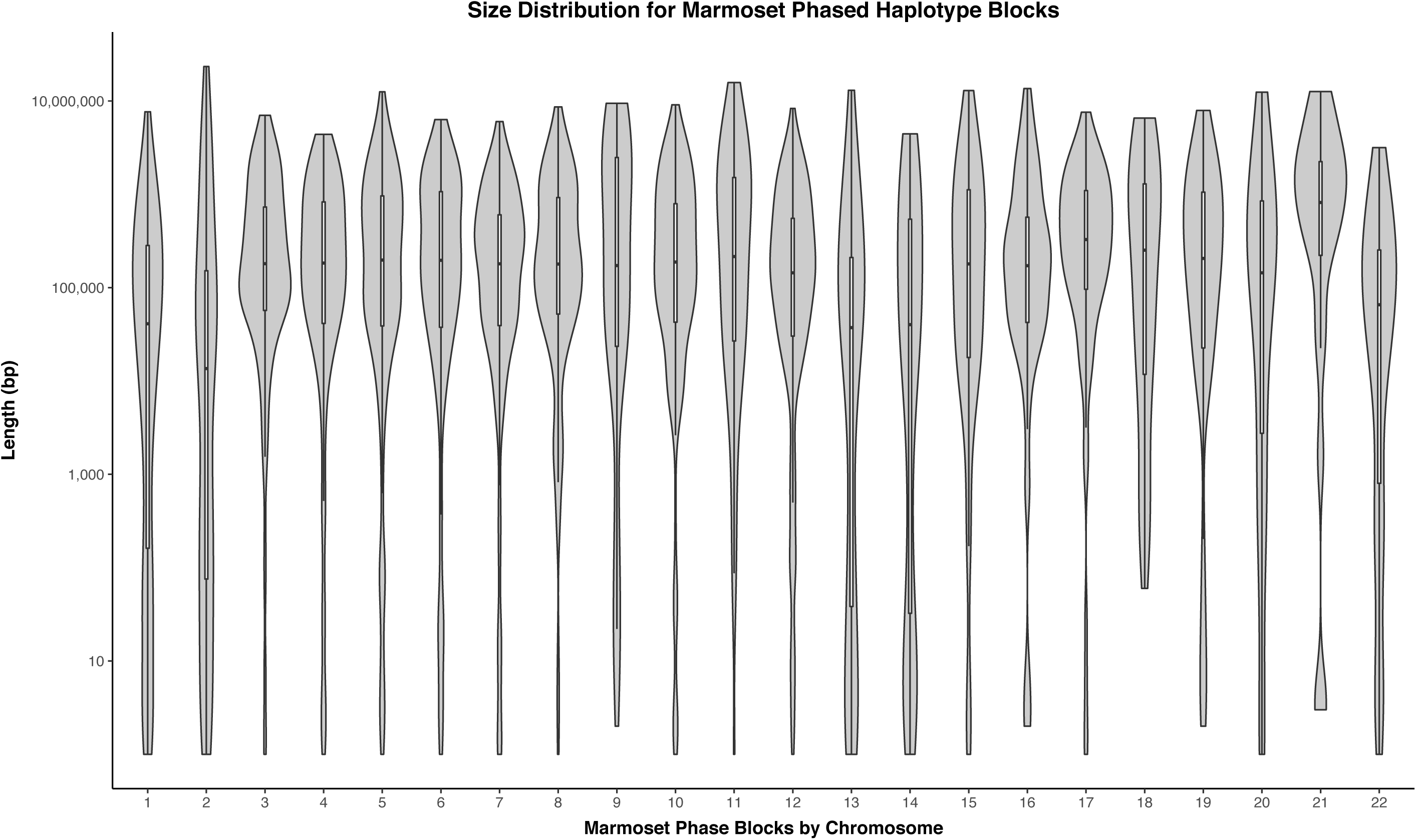
Violin plot of size distribution of haplotype blocks in cj367 by chromosomes

**Table 1**. Summary of cj367 SNVs and Indels

**Table S1**. cj367 genomic regions exhibiting continuous homozygosity.

**Table S2**. List of CRISPR sites suitable for allele-specific targeting in cj367.

## Notes

### Competing Interest Statement

The authors have declared no competing interest.

### Summary of Updates

added co-author Megan Albertelli, DVM, PhD, DACLAM Associate Professor Department of Comparative Medicine Stanford University updated supp. table 3 (added phased genotype column)

## References

Adey, A., Burton, J.N., Kitzman, J.O., Hiatt, J.B., Lewis, A.P., Martin, B.K., Qiu, R., Lee, C., and Shendure, J. (2013). The haplotype-resolved genome and epigenome of the aneuploid HeLa cancer cell line. Nature 500, 207–211.

Ausderau, K.K., Dammann, C., McManus, K., Schneider, M., Emborg, M.E., and Schultz-Darken, N. (2017). Cross-species comparison of behavioral neurodevelopmental milestones in the common marmoset monkey and human child. Dev. Psychobiol. 59, 807–821.

Benirschke, K., Anderson, J.M., and Brownhill, L.E. (1962). Marrow Chimerism in Marmosets. Science 138, 513–515.

Cai, X., Evrony, G.D., Lehmann, H.S., Elhosary, P.C., Mehta, B.K., Poduri, A., and Walsh, C.A. (2014). Single-cell, genome-wide sequencing identifies clonal somatic copy-number variation in the human brain. Cell Rep. 8, 1280–1289.

Camus, S., Ko, W.K.D., Pioli, E., and Bezard, E. (2015). Why bother using non-human primate models of cognitive disorders in translational research? Neurobiol. Learn. Mem. 124, 123–129.

Canver, M.C., Joung, J.K., and Pinello, L. (2018). Impact of Genetic Variation on CRISPR-Cas Targeting. Cris. J. 1, 159–170.

Capitanio, J.P., and Emborg, M.E. (2008). Contributions of non-human primates to neuroscience research. Lancet 371, 1126–1135.

Chen, B., Gilbert, L.A., Cimini, B.A., Schnitzbauer, J., Zhang, W., Li, G.-W., Park, J., Blackburn, E.H., Weissman, J.S., Qi, L.S., et al. (2013). Dynamic imaging of genomic loci in living human cells by an optimized CRISPR/Cas system. Cell 155, 1479–1491.

Chen, Y., Zheng, Y., Kang, Y., Yang, W., Niu, Y., Guo, X., Tu, Z., Si, C., Wang, H., Xing, R., et al. (2015). Functional disruption of the dystrophin gene in rhesus monkey using CRISPR/Cas9. Hum. Mol. Genet. 24, 3764–3774.

Debowski, K., Drummer, C., Lentes, J., Cors, M., Dressel, R., Lingner, T., Salinas-Riester, G., Fuchs, S., Sasaki, E., and Behr, R. (2016). The transcriptomes of novel marmoset monkey embryonic stem cell lines reflect distinct genomic features. Sci. Rep. 6, 29122.

DePristo, M. a, Banks, E., Poplin, R., Garimella, K. V, Maguire, J.R., Hartl, C., Philippakis, A. a, del Angel, G., Rivas, M. a, Hanna, M., et al. (2011). A framework for variation discovery and genotyping using next-generation DNA sequencing data. Nat Genet 43, 491–498.

Emborg, M.E. (2017). Nonhuman Primate Models of Neurodegenerative Disorders. ILAR J. 58, 190–201.

Farnsworth, S.L., Qiu, Z., Mishra, A., and Hornsby, P.J. (2013). Directed neural differentiation of induced pluripotent stem cells from non-human primates. Exp. Biol. Med. (Maywood). 238, 276–284.

Grow, D.A., McCarrey, J.R., and Navara, C.S. (2016). Advantages of nonhuman primates as preclinical models for evaluating stem cell-based therapies for Parkinson’s disease. Stem Cell Res. 17, 352–366.

International HapMap Consortium (2005). A haplotype map of the human genome. Nature 437, 1299–1320.

Izpisua-Belmonte, J.C., Callaway, E.M., Caddick, S.J., Churchland, P., Feng, G., Homanics, G.E., Lee, K.-F., Leopold, D.A., Miller, C.T., Mitchell, J.F., et al. (2015). Brains, genes, and primates. Neuron 86, 617–631.

Kang, Y., Zheng, B., Shen, B., Chen, Y., Wang, L., Wang, J., Niu, Y., Cui, Y., Zhou, J., Wang, H., et al. (2015). CRISPR/Cas9-mediated Dax1 knockout in the monkey recapitulates human AHC-HH. Hum. Mol. Genet. 24, 7255–7264.

Karolchik, D., Hinrichs, A.S., Furey, T.S., Roskin, K.M., Sugnet, C.W., Haussler, D., and Kent, W.J. (2004). The UCSC Table Browser data retrieval tool. Nucleic Acids Res. 32, D493–6.

Kishi, N., Sato, K., Sasaki, E., and Okano, H. (2014). Common marmoset as a new model animal for neuroscience research and genome editing technology. Dev. Growth Differ. 56, 53–62.

Knouse, K.A., Wu, J., and Amon, A. (2016). Assessment of megabase-scale somatic copy number variation using single-cell sequencing. Genome Res. 26, 376–384.

Kropp, J., Di Marzo, A., and Golos, T. (2017). Assisted reproductive technologies in the common marmoset: an integral species for developing nonhuman primate models of human diseases†. Biol. Reprod. 96, 277–287.

Kuhn, R.M., Haussler, D., and Kent, W.J. (2013). The UCSC genome browser and associated tools. Brief. Bioinform. 14, 144–161.

Langmead, B., and Salzberg, S.L. (2012). Fast gapped-read alignment with Bowtie 2. Nat. Methods 9, 357–359.

Li, H., and Durbin, R. (2009). Fast and accurate short read alignment with Burrows-Wheeler transform. Bioinformatics 25, 1754–1760.

Lorenz, R., Bernhart, S.H., Höner Zu Siederdissen, C., Tafer, H., Flamm, C., Stadler, P.F., and Hofacker, I.L. (2011). ViennaRNA Package 2.0. Algorithms Mol. Biol. 6, 26.

Luo, X., Li, M., and Su, B. (2016). Application of the genome editing tool CRISPR/Cas9 in non-human primates. Zool. Res. 37, 214–219.

Mansfield, K. (2003). Marmoset models commonly used in biomedical research. Comp. Med. 53, 383–392.

Marks, P., Garcia, S., Barrio, A.M., Belhocine, K., Bernate, J., Bharadwaj, R., Bjornson, K., Catalanotti, C., Delaney, J., Fehr, A., et al. (2018). Resolving the Full Spectrum of Human Genome Variation using Linked-Reads. Preprint at.

Marmoset Genome Sequencing and Analysis Consortium (2014). The common marmoset genome provides insight into primate biology and evolution. Nat. Genet. 46, 850–857.

Maudhoo, M.D., Ren, D., Gradnigo, J.S., Gibbs, R.M., Lubker, A.C., Moriyama, E.N., French, J.A., and Norgren, R.B. (2014). De novo assembly of the common marmoset transcriptome from NextGen mRNA sequences. Gigascience 3, 14.

McConnell, M.J., Lindberg, M.R., Brennand, K.J., Piper, J.C., Voet, T., Cowing-Zitron, C., Shumilina, S., Lasken, R.S., Vermeesch, J.R., Hall, I.M., et al. (2013). Mosaic copy number variation in human neurons. Science 342, 632–637.

McKenna, A., Hanna, M., Banks, E., Sivachenko, A., Cibulskis, K., Kernytsky, A., Garimella, K., Altshuler, D., Gabriel, S., Daly, M., et al. (2010). The Genome Analysis Toolkit: a MapReduce framework for analyzing next-generation DNA sequencing data. Genome Res. 20, 1297–1303.

Nelson, E.E., and Winslow, J.T. (2009). Non-human primates: model animals for developmental psychopathology. Neuropsychopharmacology 34, 90–105.

Nishimasu, H., Ran, F.A., Hsu, P.D., Konermann, S., Shehata, S.I., Dohmae, N., Ishitani, R., Zhang, F., and Nureki, O. (2014). Crystal structure of Cas9 in complex with guide RNA and target DNA. Cell 156, 935–949.

Niu, Y., Shen, B., Cui, Y., Chen, Y., Wang, J., Wang, L., Kang, Y., Zhao, X., Si, W., Li, W., et al. (2014). Generation of gene-modified cynomolgus monkey via Cas9/RNA-mediated gene targeting in one-cell embryos. Cell 156, 836–843.

Okano, H., Hikishima, K., Iriki, A., and Sasaki, E. (2012). The common marmoset as a novel animal model system for biomedical and neuroscience research applications. Semin. Fetal Neonatal Med. 17, 336–340.

Park, J.E., Zhang, X.F., Choi, S.-H., Okahara, J., Sasaki, E., and Silva, A.C. (2016). Generation of transgenic marmosets expressing genetically encoded calcium indicators. Sci. Rep. 6, 34931.

Parker, K.J., Garner, J.P., Oztan, O., Tarara, E.R., Li, J., Sclafani, V., Del Rosso, L.A., Chun, K., Berquist, S.W., Chez, M.G., et al. (2018). Arginine vasopressin in cerebrospinal fluid is a marker of sociality in nonhuman primates. Sci. Transl. Med.

Phillips, K.A., Bales, K.L., Capitanio, J.P., Conley, A., Czoty, P.W., ‘t Hart, B.A., Hopkins, W.D., Hu, S.-L., Miller, L.A., Nader, M.A., et al. (2014). Why primate models matter. Am. J. Primatol. 76, 801–827.

Pinello, L., Canver, M.C., Hoban, M.D., Orkin, S.H., Kohn, D.B., Bauer, D.E., and Yuan, G.-C. (2016). Analyzing CRISPR genome-editing experiments with CRISPResso. Nat. Biotechnol. 34, 695–697.

Qiu, Z., Mishra, A., Li, M., Farnsworth, S.L., Guerra, B., Lanford, R.E., and Hornsby, P.J. (2015). Marmoset induced pluripotent stem cells: Robust neural differentiation following pretreatment with dimethyl sulfoxide. Stem Cell Res. 15, 141–150.

Ran, F.A., Hsu, P.D., Wright, J., Agarwala, V., Scott, D.A., and Zhang, F. (2013). Genome engineering using the CRISPR-Cas9 system. Nat. Protoc. 8, 2281–2308.

Sasaki, E. (2015). Prospects for genetically modified non-human primate models, including the common marmoset. Neurosci. Res. 93, 110–115.

Sasaki, E., Hanazawa, K., Kurita, R., Akatsuka, A., Yoshizaki, T., Ishii, H., Tanioka, Y., Ohnishi, Y., Suemizu, H., Sugawara, A., et al. (2005). Establishment of novel embryonic stem cell lines derived from the common marmoset (Callithrix jacchus). Stem Cells 23, 1304–1313.

Sasaki, E., Suemizu, H., Shimada, A., Hanazawa, K., Oiwa, R., Kamioka, M., Tomioka, I., Sotomaru, Y., Hirakawa, R., Eto, T., et al. (2009). Generation of transgenic non-human primates with germline transmission. Nature 459, 523–527.

Sato, K., and Sasaki, E. (2018). Genetic engineering in nonhuman primates for human disease modeling. J. Hum. Genet. 63, 125–131.

Sato, K., Kuroki, Y., Kumita, W., Fujiyama, A., Toyoda, A., Kawai, J., Iriki, A., Sasaki, E., Okano, H., and Sakakibara, Y. (2015). Resequencing of the common marmoset genome improves genome assemblies and gene-coding sequence analysis. Sci. Rep. 5, 16894.

Sato, K., Oiwa, R., Kumita, W., Henry, R., Sakuma, T., Ito, R., Nozu, R., Inoue, T., Katano, I., Sato, K., et al. (2016). Generation of a Nonhuman Primate Model of Severe Combined Immunodeficiency Using Highly Efficient Genome Editing. Cell Stem Cell 19, 127–138.

Schultz-Darken, N., Braun, K.M., and Emborg, M.E. (2016). Neurobehavioral development of common marmoset monkeys. Dev. Psychobiol. 58, 141–158.

Shiozawa, S., Kawai, K., Okada, Y., Tomioka, I., Maeda, T., Kanda, A., Shinohara, H., Suemizu, H., James Okano, H., Sotomaru, Y., et al. (2011). Gene targeting and subsequent site-specific transgenesis at the β-actin (ACTB) locus in common marmoset embryonic stem cells. Stem Cells Dev. 20, 1587–1599.

Sunagawa, G.A., Sumiyama, K., Ukai-Tadenuma, M., Perrin, D., Fujishima, H., Ukai, H., Nishimura, O., Shi, S., Ohno, R.-I., Narumi, R., et al. (2016). Mammalian Reverse Genetics without Crossing Reveals Nr3a as a Short-Sleeper Gene. Cell Rep. 14, 662– 677.

Tatsuki, F., Sunagawa, G.A., Shi, S., Susaki, E.A., Yukinaga, H., Perrin, D., Sumiyama, K., Ukai-Tadenuma, M., Fujishima, H., Ohno, R., et al. (2016). Involvement of Ca(2+)-Dependent Hyperpolarization in Sleep Duration in Mammals. Neuron 90, 70–85.

The 1000 Genomes Project Consortium, Abecasis, G.R., Altshuler, D., Auton, A., Brooks, L.D., Durbin, R.M., Gibbs, R.A., Hurles, M.E., and McVean, G.A. (2010). A map of human genome variation from population-scale sequencing. Nature 467, 1061–1073.

Thomson, J.A., Kalishman, J., Golos, T.G., Durning, M., Harris, C.P., and Hearn, J.P. (1996). Pluripotent cell lines derived from common marmoset (Callithrix jacchus) blastocysts. Biol. Reprod. 55, 254–259.

Tomioka, I., Maeda, T., Shimada, H., Kawai, K., Okada, Y., Igarashi, H., Oiwa, R., Iwasaki, T., Aoki, M., Kimura, T., et al. (2010). Generating induced pluripotent stem cells from common marmoset (Callithrix jacchus) fetal liver cells using defined factors, including Lin28. Genes to Cells 15, 959–969.

Tomioka, I., Ishibashi, H., Minakawa, E.N., Motohashi, H.H., Takayama, O., Saito, Y., Popiel, H.A., Puentes, S., Owari, K., Nakatani, T., et al. (2017). Transgenic Monkey Model of the Polyglutamine Diseases Recapitulating Progressive Neurological Symptoms. Eneuro 4, ENEURO.0250-16.2017.

Vermilyea, S.C., Guthrie, S., Meyer, M., Smuga-Otto, K., Braun, K., Howden, S., Thomson, J.A., Zhang, S.-C., Emborg, M.E., and Golos, T.G. (2017). Induced Pluripotent Stem Cell-Derived Dopaminergic Neurons from Adult Common Marmoset Fibroblasts. Stem Cells Dev. 26, 1225–1235.

Vermilyea, S.C., Babinski, A., Guthrie, S., Golos, T.G., and Emborg, M.E. (2018). LRRK2 Genomic Editing in Common Marmoset Stem Cells. Cell Transplant. 27, 682– 722.

Wise, S.P. (2008). Forward frontal fields: phylogeny and fundamental function. Trends Neurosci. 31, 599–608.

Zahs, K.R., and Ashe, K.H. (2010). “Too much good news” - are Alzheimer mouse models trying to tell us how to prevent, not cure, Alzheimer’s disease? Trends Neurosci. 33, 381–389.

Zhang, Y., Pak, C., Han, Y., Ahlenius, H., Zhang, Z., Chanda, S., Marro, S., Patzke, C., Acuna, C., Covy, J., et al. (2013). Rapid single-step induction of functional neurons from human pluripotent stem cells. Neuron 78, 785–798.

Zheng, G.X.Y., Lau, B.T., Schnall-Levin, M., Jarosz, M., Bell, J.M., Hindson, C.M., Kyriazopoulou-Panagiotopoulou, S., Masquelier, D.A., Merrill, L., Terry, J.M., et al. (2016). Haplotyping germline and cancer genomes with high-throughput linked-read sequencing. Nat. Biotechnol. 34, 303–311.

Zhou, B., Ho, S.S., Zhu, X., Zhang, X., Spies, N., Byeon, S., Arthur, J.G., Pattni, R., Ben-Efraim, N., Haney, M.S., et al. (2018). Comprehensive, integrated and phased whole-genome analysis of the primary ENCODE cell line K562. bioRxiv.

